# Contribution of the axon initial segment to action potentials recorded extracellularly

**DOI:** 10.1101/243808

**Authors:** Maria Teleńczuk, Romain Brette, Alain Destexhe, Bartosz Teleńczuk

## Abstract

Action potentials (APs) are electric phenomena that are recorded both intracellularly and extracellularly. APs are usually initiated in the short segment of the axon called the axon initial segment (AIS). It was recently proposed that at onset of an AP the soma and the AIS form a dipole. We study the extracellular signature (the extracellular action potential, EAP) generated by such a dipole. First, we demonstrate the formation of the dipole and its extracellular signature in detailed morphological models of a reconstructed pyramidal neuron. Then, we study the EAP waveform and its spatial dependence in models with axonal AP initiation and contrast it with the EAP obtained in models with somatic AP initiation. We show that in the models with axonal AP initiation the dipole forms between somatodendritic compartments and the AIS, and not between soma and dendrites as in the classical models. Soma-dendrites dipole is present only in models with somatic AP initiation. Our study has consequences for interpreting extracellular recordings of single-neuron activity and determining electrophysiological neuron types, but also for better understanding the origins of the high-frequency macroscopic electric fields recorded in the brain.

**New & Noteworthy:** We studied the consequences of the action potential (AP) initiation site on the extracellular signatures of APs. We show that: (1) at the time of AP initiation the action initial segment (AIS) forms a dipole with the soma, (2) the width but not (3) amplitude of the extracellular AP generated by this dipole increases with the soma-AIS distance. This may help to monitor dynamic changes in the AIS position in experimental in vivo recordings.

## Introduction

Action potentials (APs) are the main output of neuronal computation arising due to neuronal membrane excitability. The most direct method to detect APs is by intracellular recordings for which a glass pipette is inserted into the soma. However, the sample size of neurons recorded with this technique is limited. Another method of AP detection uses extracellular electrodes whose densities can be greatly increased thanks to the silicon technology opening the possibility of massive recordings from large samples of neurons (Jun et al., 2017; Stevenson and Kording, 2011). The drawback of this method is that the discrimination of separate neurons and their types based on extracellular recordings is not trivial (Barthó et al., 2004) and requires a detailed model of how the extracellular signature of the APs is generated.

APs also contribute to the local field potentials (LFP) and electroencephalograms (EEG) recorded far from the neuronal source. In particular, the high-frequency components of these signals can relate to the firing rates of large population of neurons (Reimann et al., 2013). These high-frequency local field potentials are also known to be sensitive to the neuronal responses at single-neuron and single-trial level (Telenczuk et al., 2015). Therefore, APs can be as important as the passive dendritic and synaptic currents for understanding the LFP or EEG and in particular their high-frequency components.

The extracellular signature of APs has been a topic of computational studies (Bédard et al., 2004; Gold et al., 2006; Milstein and Koch, 2008). These studies emphasize the role of passive currents and dendritic compartments in the generation of the action potentials. However, in most of those models APs were initiated in the soma. It is now well established that the AP often initiates in the axon initial segment (AIS) (Stuart et al., 1997a; Stuart et al., 1997b), which gives a characteristic kink at the AP onset when recorded somatically (Naundorf et al., 2007). This kink can be explained by the “critical resistive coupling model”, according to which the AP is initiated through the strong resistive coupling between a small AIS and a large soma (Brette, 2013; Telenczuk et al., 2017). In this mechanism of AP initiation, AIS and soma form effectively a current dipole.

We studied the contribution of the soma-AIS dipole to the extracellular field and its effect on the shape and amplitude of the extracellular action potential (EAP). In particular, we studied the EAP from realistic model neurons with AIS-based initiation and compare it with models for which the sodium channel density was modified to initiate the AP somatically. By means of computational modelling, we show that the AIS contributes significantly to the EAP. Although, the localization and length of the AIS have only a minor effect on the appearance of the AP recorded intracellularly from the soma, the presence of AIS has a large impact on the shape of the EAP.

We believe that these findings improve our understanding of the close-field and far-field contribution of the AP to the electric fields in the brain. It will also help to interpret recordings of various signals ranging from the EAP, through LFP to EEG.

## Materials and Methods

### Detailed morphology model

We used a detailed morphology model (physiological Nav model) of the rat neocortex, layer 5 pyramidal neuron described in Hallermann et al. (2012), whose morphology and ion channels are modelled such as to give good fit to the experimental data. Most importantly, in this model action potentials initiate in the axon initial segment as is the case in real neurons. The details of the model can be found in (Hallermann et al., 2012).

The kinetics of the sodium channels were matched to experimental data recorded from the soma (putative Nav1.2 channel). Another sodium channel (putative Nav1.6 channel) was introduced in the axon, of which activation curve was shifted by 2 mV towards more negative potential to account for the lower threshold of AP initiation in the AIS.

The density of the sodium channels in the soma was 500 *pSμm*^−2^ while in the AIS it varied between 1452 and 8392 *pSμm*^−2^ (Fig. 2C physiological Nav model). To compare the results of the original model to the neuron where the action potential initiates in the soma, we reduced the density of the sodium channels in the AIS just below that in the soma to 480 *pSμm*^−2^ throughout the length of the initial segment of the axon (70 *μm*, Fig. 2C, reduced Nav model). The density of the sodium channels in the soma remained the same as in the original model (500 *pSμm*^−2^). This was enough for an action potential to initiate in the soma. We note that there are less sodium channels in the altered model leading to lower current flow, therefore comparison of the absolute amplitudes of the extracellular potential is not possible. Therefore, where necessary, we normalized the potentials to the highest absolute value of the potential (Fig. 5 and 6).

To trigger the action potential we injected current to the soma. To remove signal associated with the current injection we removed all the active channels from the model and stimulated it in the same way. We then subtracted the results of the passive model from the results of each of the active models.

In the detailed morphology model, in all the calculations, the soma is represented as a cylinder. However, in the figures we represent it as a triangular shape for easier visualisation of the morphology of the cell.

### Soma-axon model

We used a simple neuron consisting of a soma (20 × 30 *μ*m, 6 segments) and an axon (1 × 50 *μ*m, 10 segments), adapted from (Yu et al., 2008). Figure 8A shows the sample schematics of the shape of the neuron. The simulation was controlled from Python using the Neuron-Python interface (Hines et al., 2009).

### Linear Source Approximation

To estimate the extracellular potential, we used the Linear Source Approximation (LSA) method, which calculates the summed potential generated by currents originating from line sources with known sizes and positions. This method is known to be more precise than approximating the currents by point sink and sources (Holt, 1997; Wilson and Bower, 1992). We then applied the LSA estimation to cylinders obtained from the segmentation by Neuron simulator (Hines and Carnevale, 1997). The field was calculated using the LSA implementation of NeuronEAP Python library (Telenczuk and Telenczuk, 2016). In all calculations we used an extracellular conductivity of 0.3 Sm^−1^ (Nunez and Srinivasan, 2006).

In Figure 8 we removed the baseline from the extracellular potential by calculating an average potential in a window of 2 to 1 ms before the peak of the action potential.

## Results

### AP is initiated in the AIS and gives a characteristic “kink” to the somatic potential

To determine the contribution of an AP to the electric field recorded around the neuron, we performed simulations of a detailed reconstruction of a thick-tufted pyramidal neuron (neocortex, layer 5, rat, Fig. 1). The morphology reflected real reconstructed neurons with all neuronal compartments including an axon and dendrites. The densities and the kinetics of sodium (Na) and potassium (K) channels in soma and axon were constrained by the experimental data. In particular, two different types of sodium channels were introduced (referred to as Nav1.2 and Nav1.6, see Methods) with different voltage activation threshold and different distribution of the channel density across the axosomatic axis (Fig. 2, left). Overall, this model has been found to match well the properties of AP initiation in cortical neurons (Hallermann et al., 2012; Telenczuk et al., 2017).

**Figure 1:**
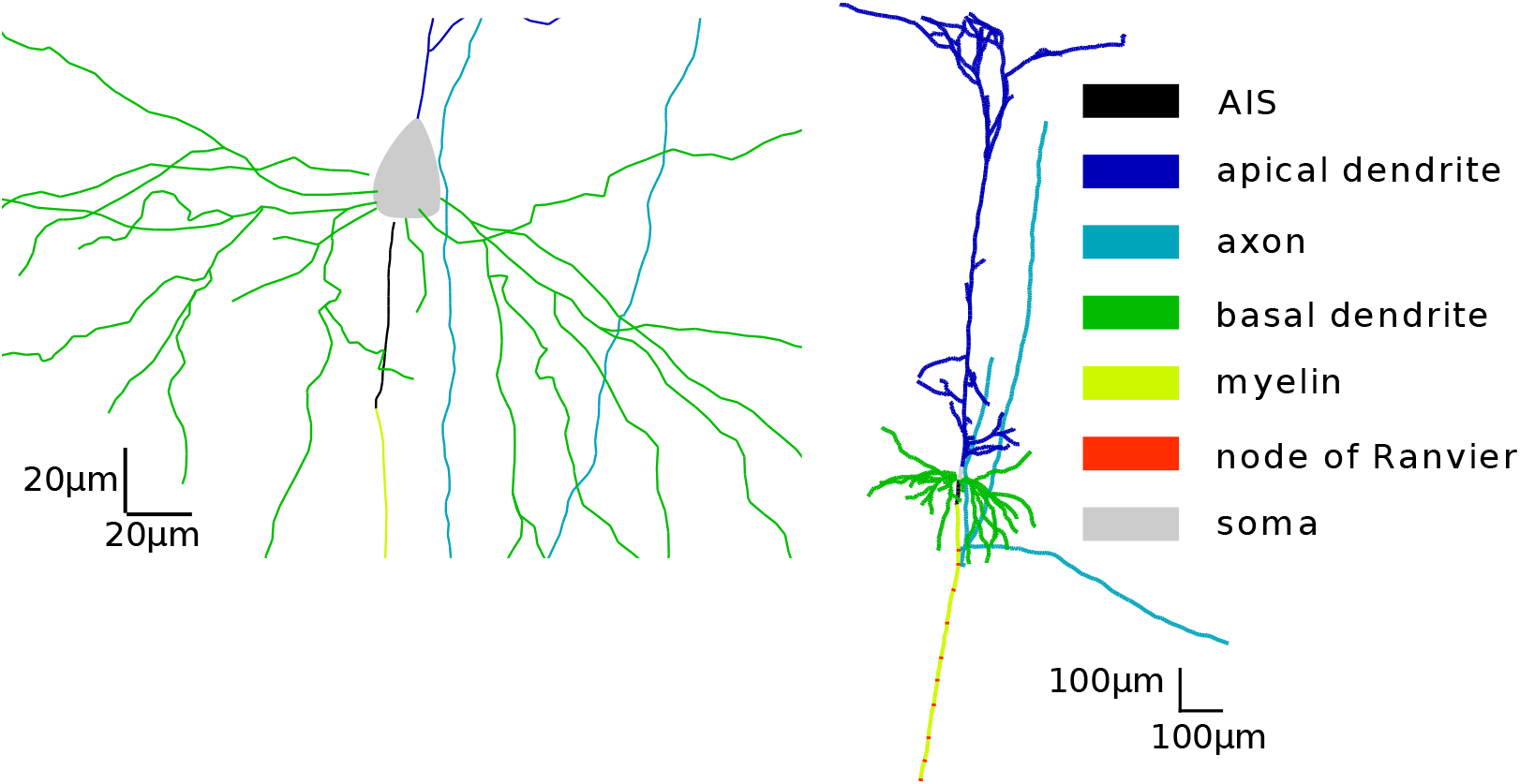
Morphology of the full compartmental model. Left: zoom into the AIS

**Figure 2.:**
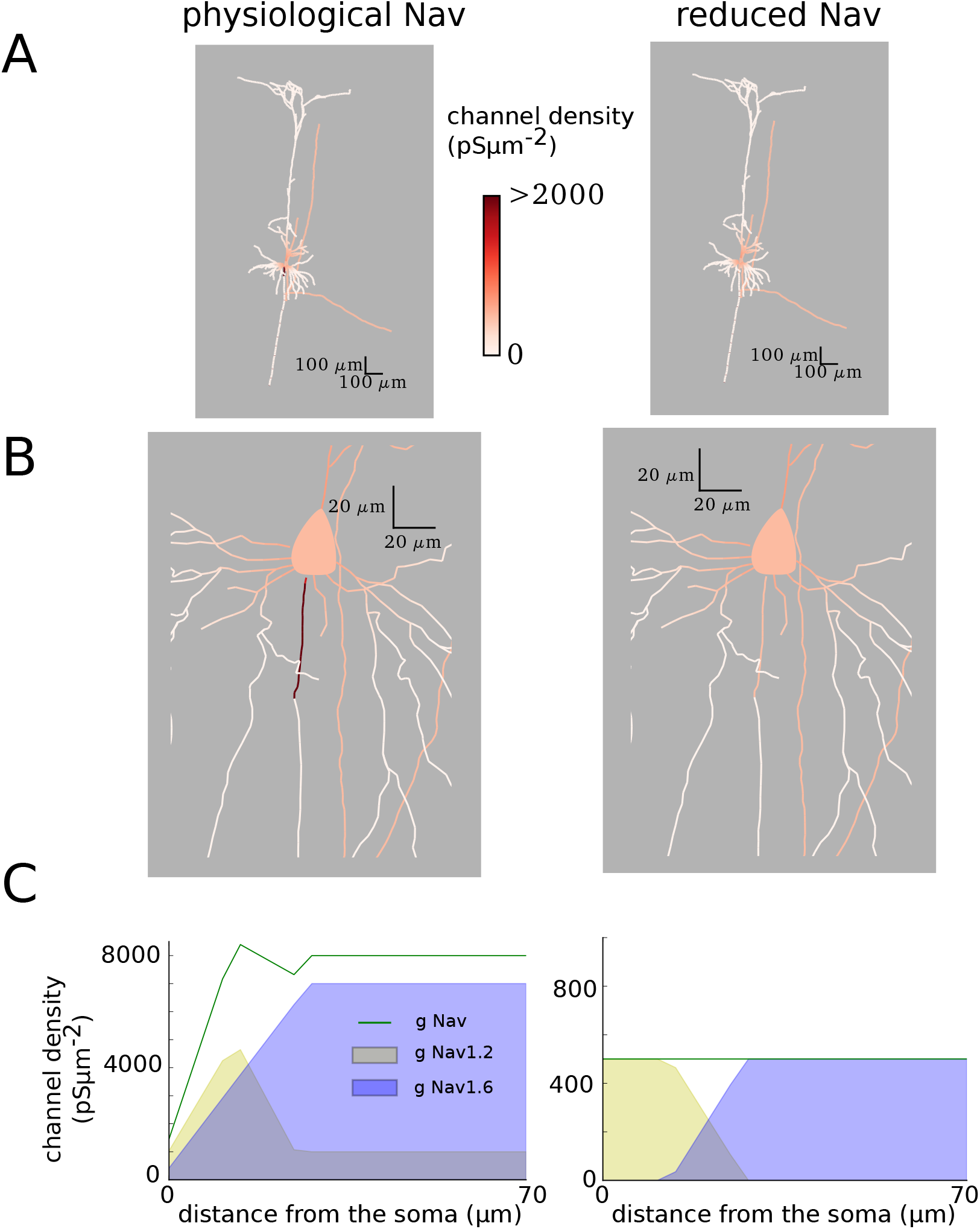
Sodium distribution within the neuron. Color scale shows the channel conductance per membrane area. *Left:* Physiological Nav model, *Right:* Reduced Nav model. *A:* Full morphology. *B:* Zoom in into the soma and the initial segments of the axon. *C* : Concentrations of two different types of sodium channels (Nav1.2 and Nav1.6) in the AIS (at 0 *μm* AIS is attached to the soma, 69.90 *μm* is its far end). Note that in both models, the density of Nav1.2 channels in the soma is 500 *pSμm*^−2^ while there are no Nav1.6 channels.

Importantly, in this model the action potential initiates distally from the soma, in the axon initial segment (AIS), and later triggers a somatic AP which is in agreement with physiological recordings (Stuart et al., 1997a). This mechanism of AP initiation gives a characteristic “kink” at the onset of the somatic AP (Fig. 3A). This is consistent with resistive coupling between the AIS and soma (Telenczuk et al., 2017). The resistive coupling model predicts that the soma and AIS form a dipole at AP initiation, which should be observed in the extracellular electric field.

**Figure 3:**
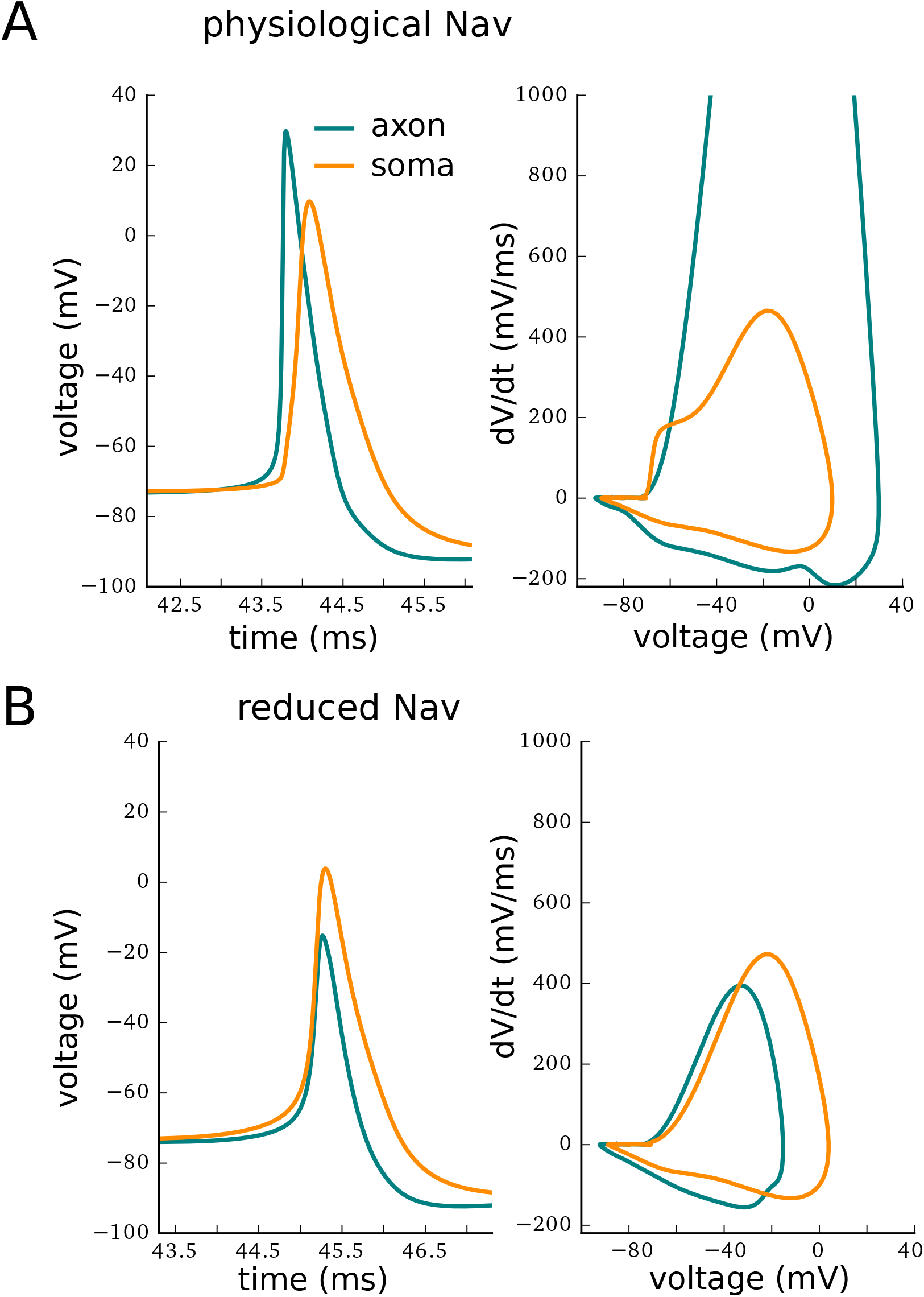
Action potentials in two different locations: soma (orange) and AIS (blue). The AP is shown both in time domain (left) and in a phase-plot (right). *A*: Physiological Nav model. *B*: Reduced Nav model.

### AIS generates positive peak at the onset of the EAP

We first characterised the waveform of the extracellular action potential (EAP). Previous models displaying somatic AP initiation have indicated that mainly sodium currents in the soma and dendrites might contribute to the initial phases of the EAP, whereas later phases are shaped by the repolarisation mediated by potassium currents in these compartments (Gold et al., 2006). In contrast, in these models axon, distal dendrites and the capacitive current contribute little to the EAP.

We re-evaluated the contribution of the AP to the extracellular potential in the more realistic model with AIS-initiated AP. First, we calculated and plotted the EAP recorded in the perisomatic area covering soma, proximal dendrites and the AIS in the physiological Nav model (Fig. 4). Consistently with previous results (Gold et al., 2006), we found a large and sharp negative peak, due to sodium inflow, followed by a broad positive peak, due to potassium-based repolarisation of the soma and dendrites. Interestingly, in some electrodes (around and above soma) these peaks were preceded by a sharp positive deflection reflecting strong axial currents flowing between AIS and soma at the onset of the AP.

**Figure 4:**
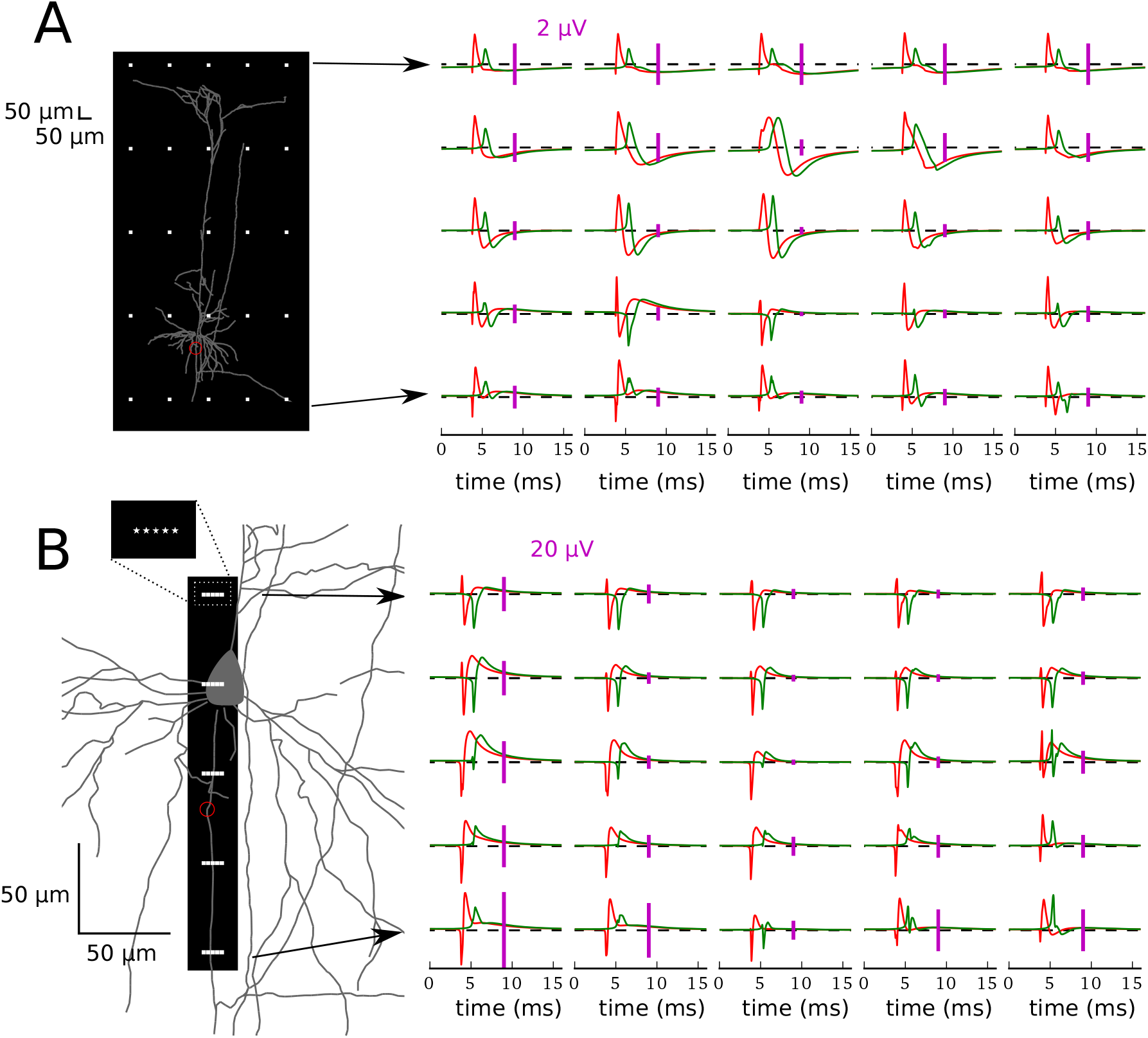
Extracellular potential (right) measured at different locations (white dots within the black rectangle, left) for the physiological Nav model (red) and reduced Nav model (green). Scale bars (pink) of 2 *μV* (A) and 20 *μV* (B) are shown for each panel separately. The y-scale is adjusted in each panel separately for better visualisation of the EAPs. **A**, Full morphology and **B**, Zoom in to the soma and initial part of the axon. The distal end of the AIS is marked with a red circle (A-B).

To confirm that this initial positive peak is related to the resistive coupling between soma and AIS forming a dipole, we lowered the densities of sodium channels in the AIS (Fig. 2, right). As expected, this modification led to the somatic initiation of the AP, which appears simultaneously at soma and AIS (these two compartments being almost isopotential), and longer AP latency due to higher threshold (Fig. 3). The EAP waveforms obtained in this modified model lack the initial positivity consistently with the results of Gold et al. (2006). We emphasise though that such a model is inconsistent with the experimental observations of AP initiation, which support axonal (AIS) rather than somatic initiation of APs.

### The AIS enhances the EAP amplitude at broad spatial ranges

The peak-to-peak amplitude decays with the distance from the neuron (Fig. 5). It is highest around soma and AIS, where the largest inflow of sodium and outflow of potassium during the AP takes place. Lowering sodium channel density such that AP initiates somatically attenuates the peak-to-peak amplitude of the EAP, which is expected from the decrease of the total membrane current in the low-sodium model (not shown). Importantly, the reduction of EAP amplitude was most pronounced in the axonal region, especially in the proximity of the axon segment previously acting as the AIS (Fig. 6).

**Figure 5.:**
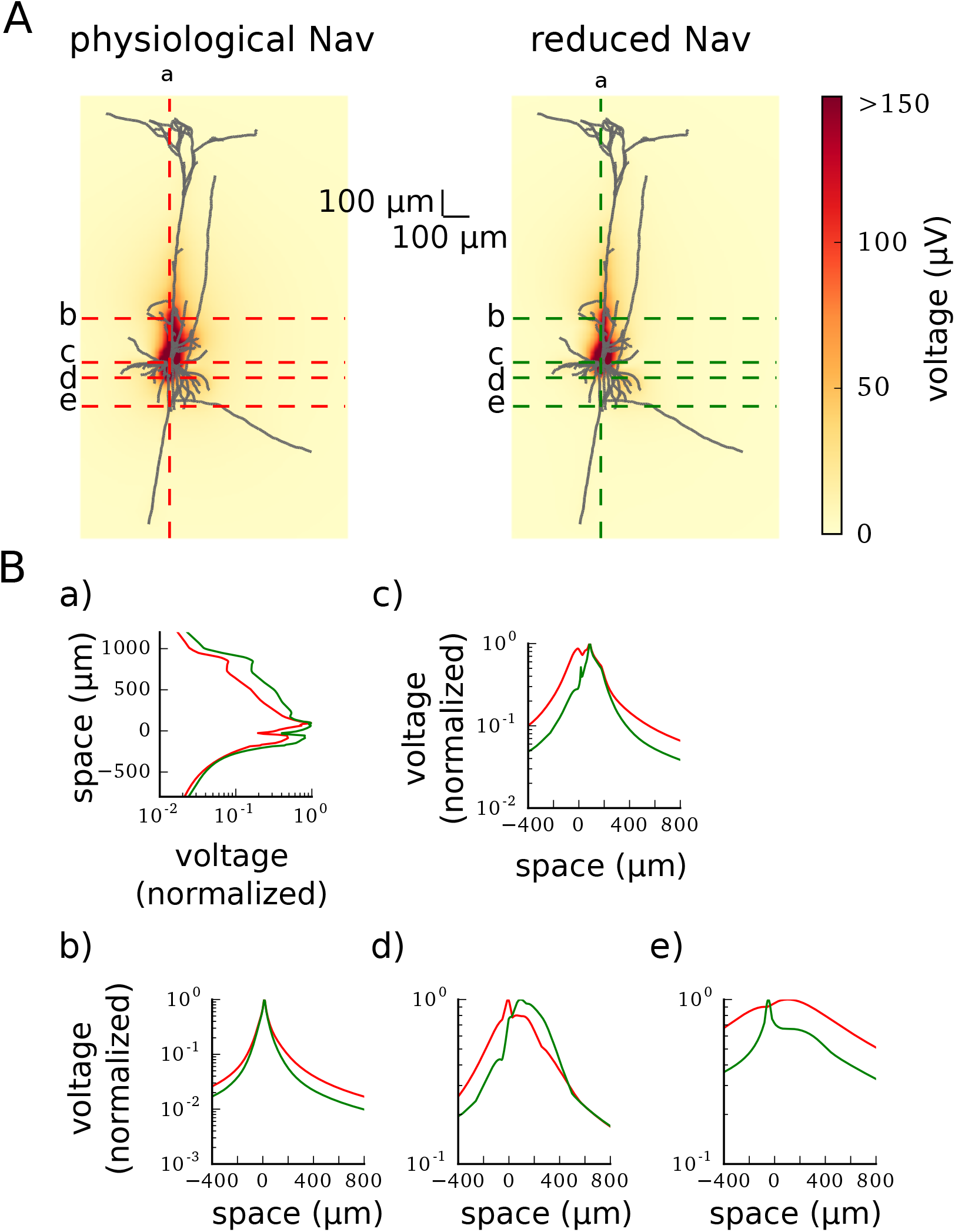
Maximum peak-to-peak amplitude of the EAP calculated in the different places of the field. *A*: Full morphology imposed on the maximum EAP amplitude (heatmap, colorbar on the right) in the physiological Nav model (left) and reduced Nav model (right). The highestamplitude EAPs are obtained in the somatic region of the neuron (dark red color in heat map, see also Figure 6 for a zoom-in). Dotted lines show the axes along which sub-panels (a)-(e) of (B) are calculated. Soma is centered at the position (0 *μm,* 0 *μm*). *B*: Maximum peak-to-peak potential normalized to the largest value of the potential for each model separately. The potential is given in the logarithmic scale. (a) Signal recorded in the vertical axis passing through the soma, (b) signal recorded in the horizontal axis passing 200 *μm* above the soma, (c) signal recorded in the horizontal axis passing through the soma, (d) signal recorded in the horizontal axis passing through the AIS, (e) signal recorded in the horizontal axis passing through 200 *μm* below the soma.

**Figure 6:**
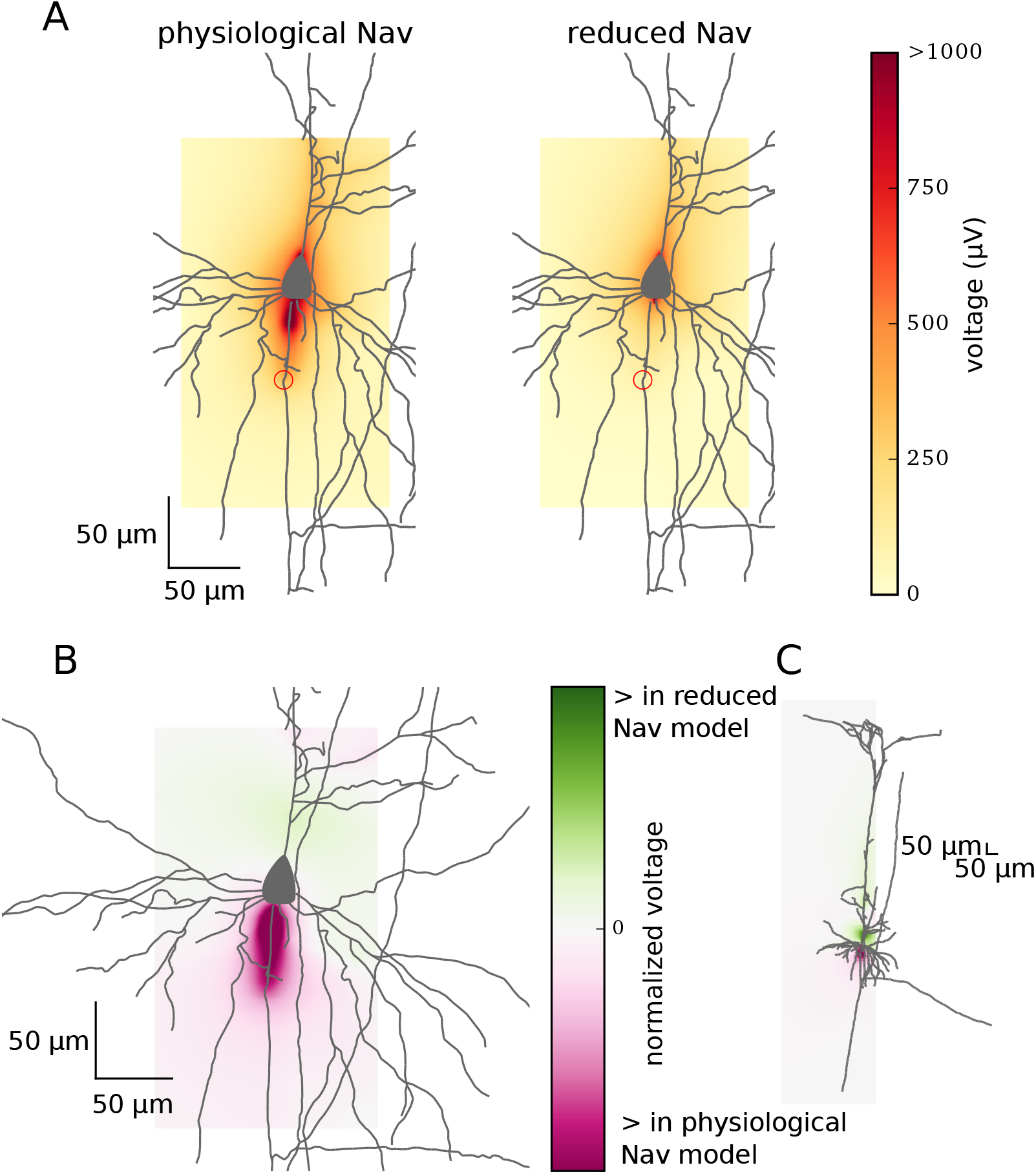
Comparison of EAP amplitude in physiological and reduced Nav models. *A:* Zoom in to the maximum peak-to-peak amplitude of the EAP (shown as heatmap, colorbar is on the right-hand side) generated by the physiological Nav model (left) and reduced Nav model (right). The amplitude around AIS (red circles – distal end) is higher in the model with axonal initiation (physiological Nav model). *B-C*: Difference between normalized peak-to-peak amplitudes (heatmap, colobar on the right) of the EAP obtained from physiological and reduced Nav models: the zoomed in view (B) and full morphology (C).

Next, we plotted the peak-to-peak amplitude of the EAP across four lines perpendicular to the somatodendritic axis (Fig. 5). Close to the neuron the profile of the EAP amplitude was non-monotonic due to the complex morphology of the neuron but it monotonically decreased with distance further away from the source. Again, due to the larger total membrane current the EAP amplitude is greater in standard sodium models compared to the low-sodium modification across all distances.

### AIS contribution to EAP can be approximated by a soma-AIS dipole

In the physiological Nav model, at the moment of AP initiation the axial current and the extracellular currents form a current loop. This current loop produces extracellular potential with dipolar configuration, i.e. negative potential around AIS (sink) and positive potential around soma and proximal dendrites (source, Fig. 7B). This relation is reversed during the repolarisation phase of the AP during which the polarities of AIS and somatodendritic compartments are reversed (Fig. 7C). Such a configuration of sinks and sources will be referred to as soma-AIS dipole. In the model with somatic AP initiation (reduced Nav model), the soma and AIS are almost isopotential so no current flows between them. In this case the soma-AIS dipole is not formed, but it is replaced by the source in the soma (or sink after the inversion) and the sink in proximal dendritic tree (soma-dendrites dipole)

**Figure 7:**
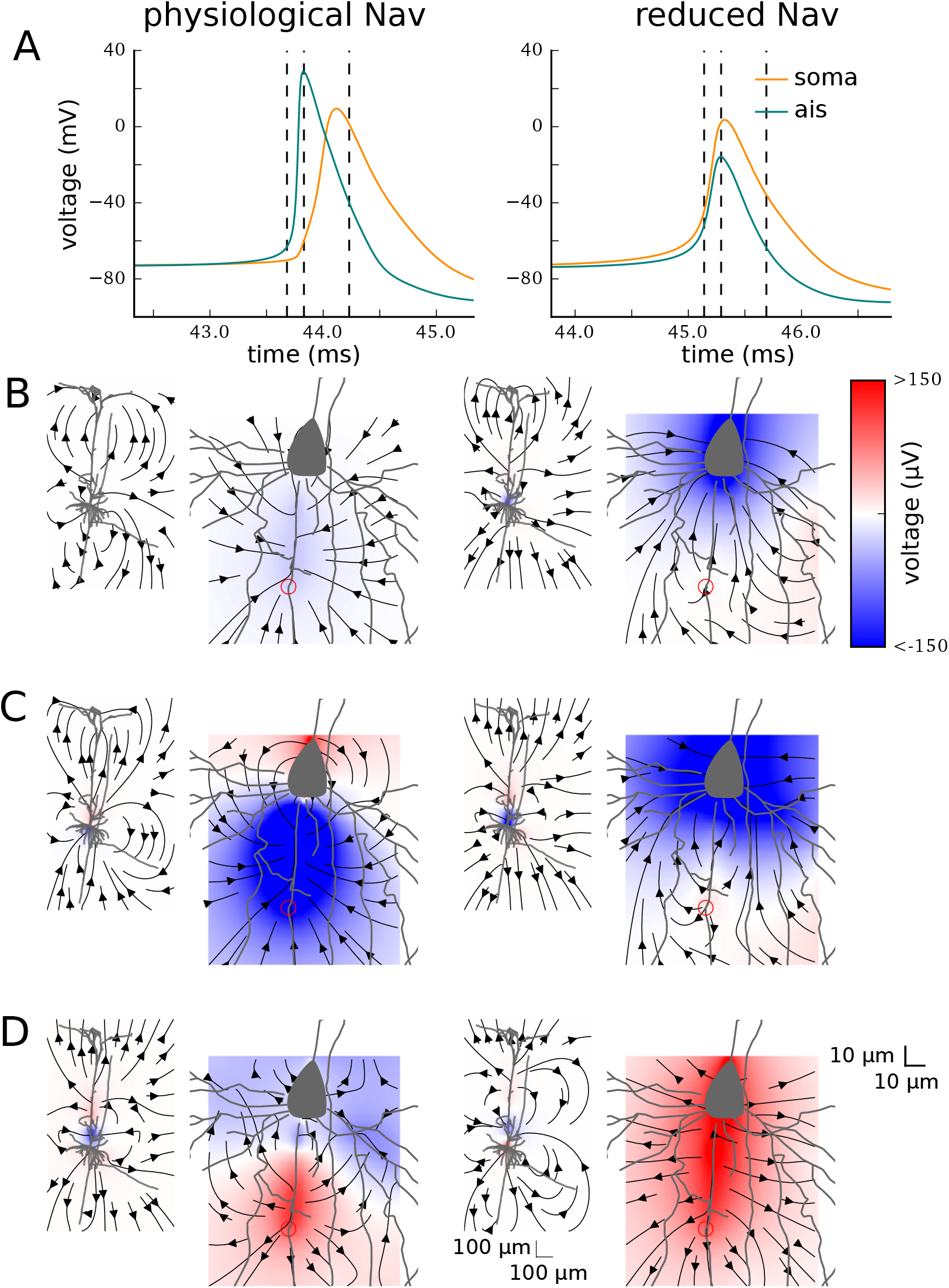
EAP at different time points in the physiological Nav model (left) and reduced Nav model (right). A, Intracellular APs in the soma (orange) and in the end of the AIS (blue). Dotted vertical lines show at which time points B-D are recorded. B–D, Extracellular potential (colormap, see the colorbar on the right, red is positive and blue is negative) and electrical current (arrows) at different times of APs plotted for around whole morphology (left) and around the soma-AIS region (right). Recordings were made at: 0.15 ms before the peak of the AP in the AIS (B), at the peak of the AP in the AIS (C), 0.4 ms after the peak of the AP in the AIS (D). In the physiological Nav model the AP initiates in the AIS (red circles) giving rise to a dipolar potential (AIS-negative, soma-positive; C, left), which later reverses in polarity (AIS-positive, soma-negative; D, left). In contrast, reduced Nav model produces a large dipole that encompasses the axon, soma and proximal dendrites (soma-dendrites dipole).

The electric field obtained from the detailed morphological models contain a mixture of contributions from passive dendritic compartments and active axonal/somatic compartments giving rise to a complex configuration of current sinks and sources. To isolate the effects of the soma-AIS dipole and its contribution to the far-field potential, we decided to further corroborate the consequences of the “critical resistive coupling” with a simplified electric dipole model. We reduced the model to a cylindrical soma and an axon. All Nav and K channels were placed in the AIS modelled as a 5-*μ*m-long segment of the axon located 45 *μ*m distally from the soma. We have shown previously (Telenczuk et al., 2017) that this model approximates well the dipolar field also observed in the detailed morphological model described above (Fig. 7).

We calculated the extracellular potential generated by this model neuron along a line that extended from the soma-AIS axis (Fig. 8A). The amplitude of the EAP decayed monotonically with the distance from the soma (Fig. 8B). We repeated the calculation for three different distances of AIS from the soma (0, 20 and 45 *μ*m), in all cases we saw similar decay with the recording distance; the absolute amplitudes of EAP depended only slightly on the AIS position (color lines in Fig. 8B). To determine the law of EAP amplitude decay, we fitted a linear function in double logarithmic scale (i.e. both the amplitude and recording distance, *r*, were log-transformed). The slope of this function provided the estimate of the power law scaling (*k* in *r^k^* relation). We found that the EAP amplitude decayed with the inverse square of the distance from the soma (*k* ≈ –2, Fig. 8C). This inverse-square law is theoretically predicted by a dipole, when the distance to the dipole is much greater that the separation between the current source and sink (far-field approximation, Fig. 9) (Griffiths, 1999; Nunez and Cutillo, 1995). Note also that the profile of the potential obtained in detailed morphology models did not agree with this prediction. As discussed above the potential in these models changes non-monotonically with the distance from soma (Fig. 5B), likely due to the contribution of dendritic compartments dominating EAP at low frequencies.

**Figure 8:**
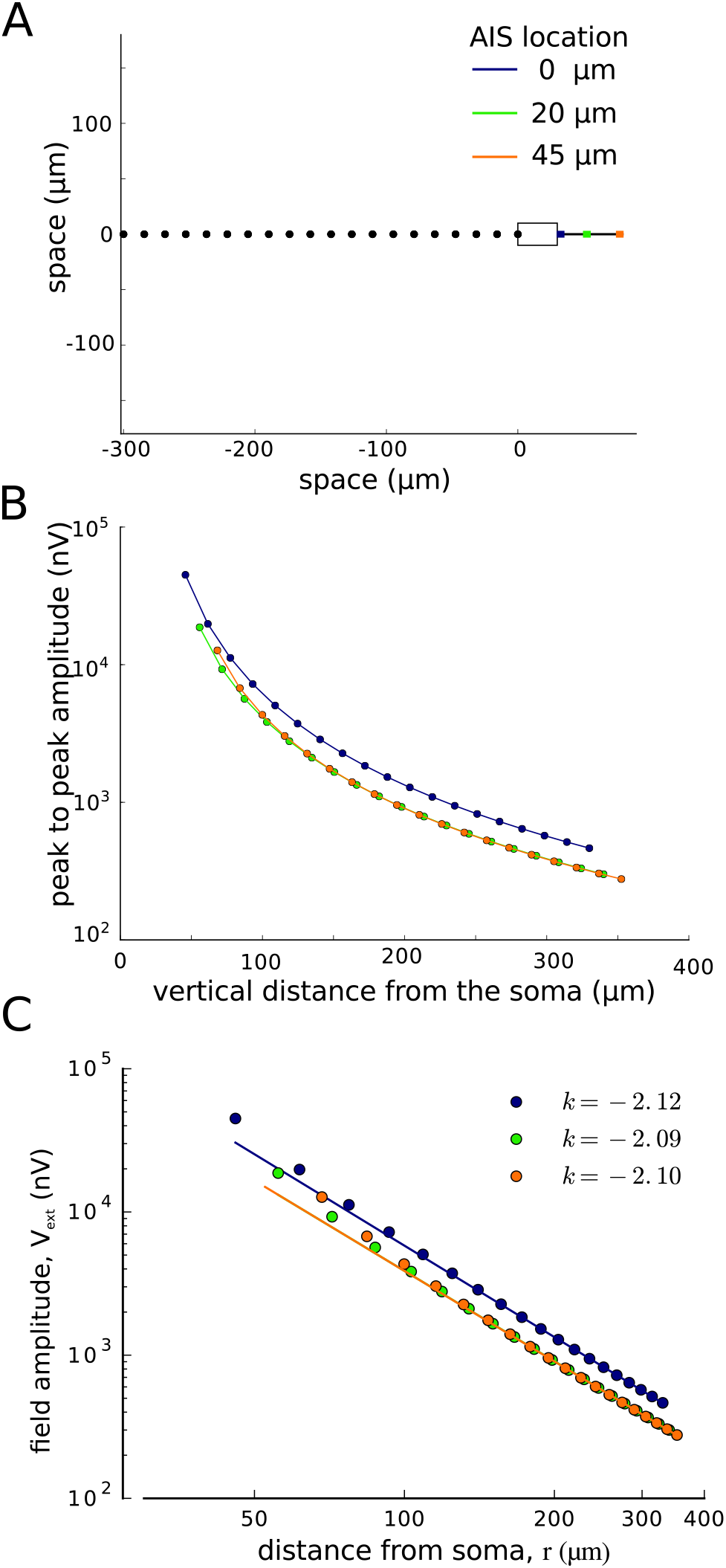
Extracellular potential calculated from the somaaxon model with the AIS at three different positions: 0 *μm* from the soma (blue), 20 *μ*m from the soma (green) and 45 *μ*m from the soma (orange). *A:* Each dot represents the location of the measurement horizontally from the soma. Schematics shows the cell body (left) and the axon (grey) with the AIS at different locations (color-coded). *B*: Logarithmic plot of the peak-to-valley amplitude of the extracellular potential vs the distance of the recording site from the soma. Color lines correspond to different positions of the AIS (see color code in A). *C*: The decay of far-field potential with distance is well approximated with a power law, *r^k^*. The exponent, *k*, estimated from the slope of linear fit to the log-transformed potential and distance is close to -2 (the value of *k* estimated for each model is given in the legend).

**Figure 9:**
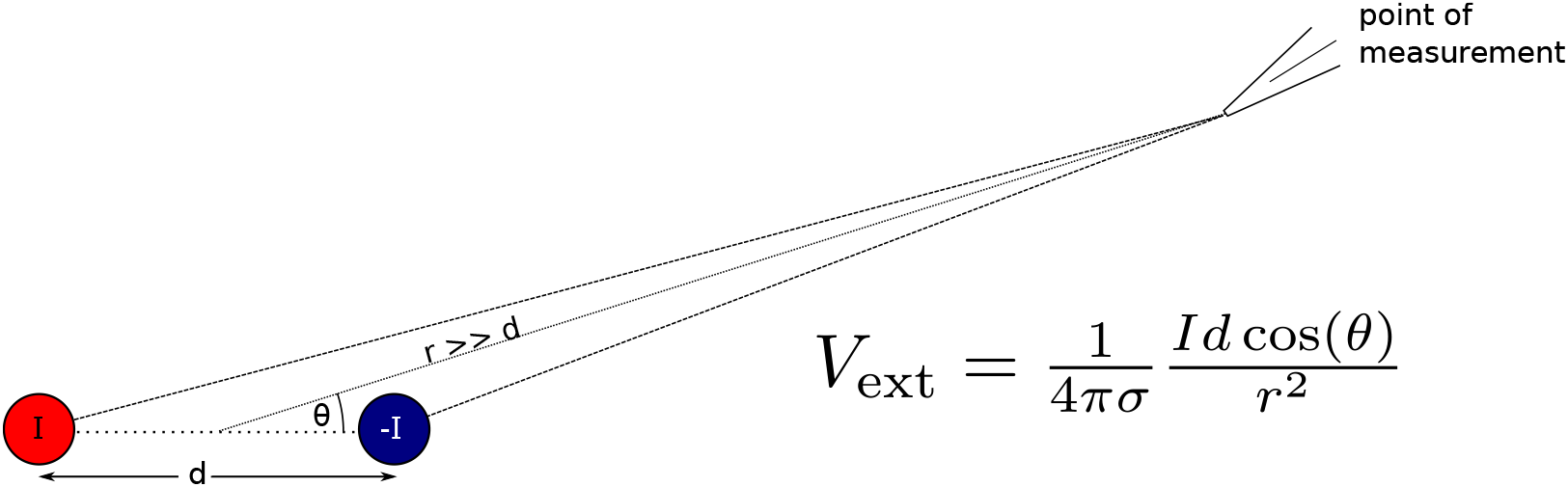
Dipole model consisting of a current sink (red) and a current source (blue) seperated by *d*. Point of measurement represents a possible recording location where extracellular potential *V*_ext_ is recorded. For the farfield approximation to hold the distance from the dipole *r* should be much larger than the distance between the sink and source (*d*). See text for more detail. *I* is current intensity, *σ* is extracellular medium conductivity, and *θ* is the angle measured from the dipole axis.

### EAP amplitude weakly depends on the distance of AIS from soma

We next investigated whether the AIS position can influence the amplitude of the EAP. The amplitude of the far-field dipole potential measured at fixed position depends on the product between the dipole current (*I*, axial current between soma and AIS) and separation between the poles (*d*, the distance from soma to the AIS; Fig. 9). Therefore, increasing the distance of the AIS from soma might increase the amplitude of the EAP, but numerical simulations of the simplified soma/AIS model showed only weak dependence of the EAP amplitude on the AIS position (Fig. 8).

To explain this finding, we investigated the effect of the AIS position on the axial current generated during the action potential. We found that the amplitude of the axial current decreased with the inverse of the distance between the AIS and the soma, *l* (Fig. 10A). Indeed, we found that it was possible to fit a straight line of slope *a* = – 1 through the points representing the logarithm of the maximum axial current versus the logarithm of the soma-AIS distance (Fig. 10B). This linear relation confirms that the amplitude of the axial current is inversely proportional to the distance between the soma and the AIS, *I_axial_* ∼ 1/*l*. Such a relationship is also predicted by the resistive coupling hypothesis (Hamada et al., 2016). This drop of current magnitude compensates for the increase between the sink and source of the dipole (soma and AIS). Since the product of current intensity, *I*, and the dipole dimension, *d*, remains constant, the EAP amplitude does not depend on the AIS position.

**Figure 10:**
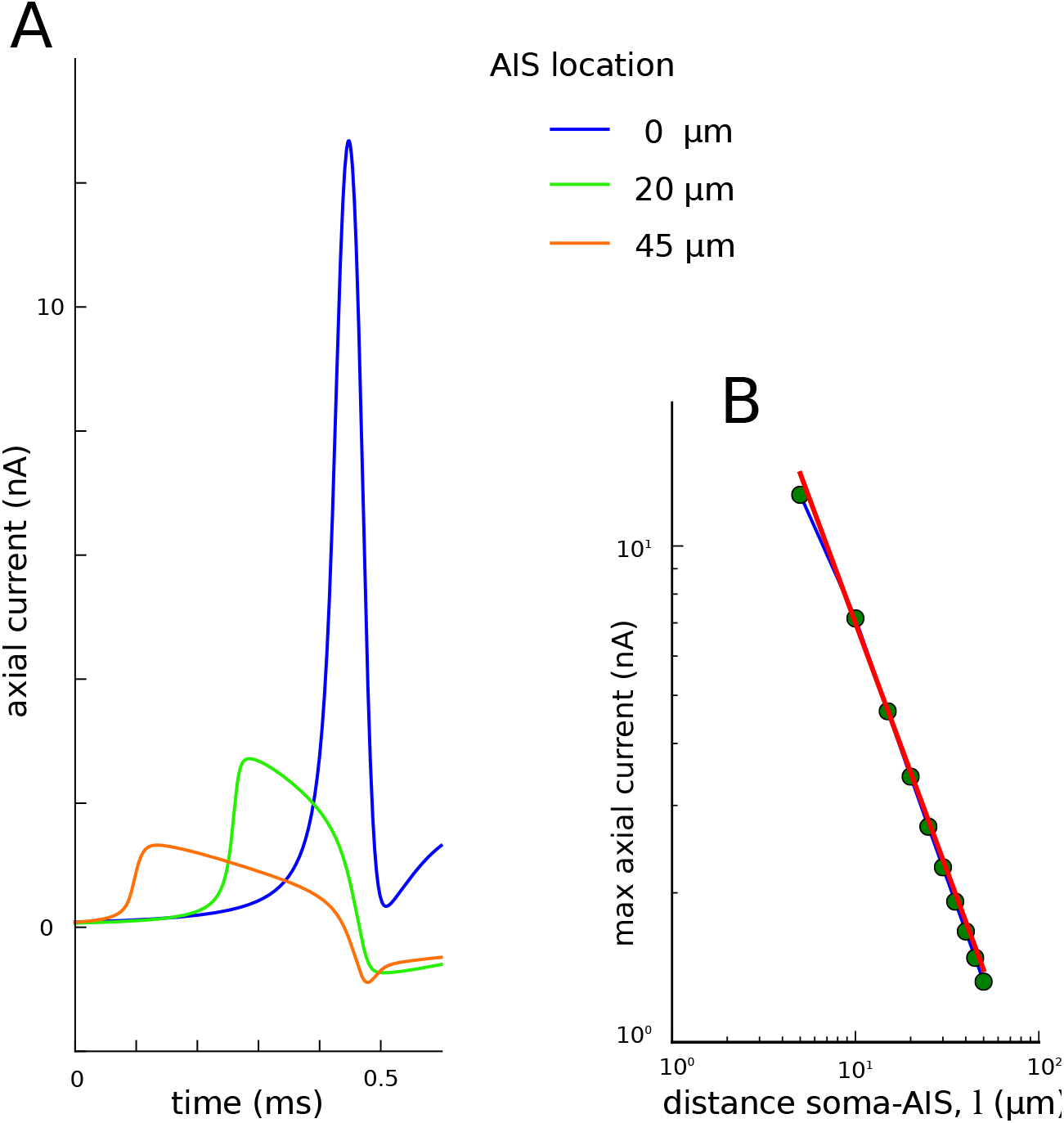
Dependence of axial current amplitude on the distance between the soma and the AIS in the soma-axon model. *A*: Axial current passing from the axon to the soma during the action potential, aligned to the peak of somatic AP (which is at 0.5 ms). *B*: The maximum of axial current vs the distance of the AIS end proximal to the soma in double-logarithmic scale. Red line shows the fitted function *I_axial_* = (70 nA · *μm*)/*l* (which is a linear function in double-logarithmic scale).

### EAP broadens with AIS distance from soma

To study the effect of the AIS position on the EAP width, we calculated the extracellular potential generated by models with the AIS placed at ten different positions from the end of the soma, up to 45 *μm* distally. We observed that the EAPs become gradually wider with increasing distance between the soma and the AIS (Fig. 11B), while the shapes of intracellular waveforms remain similar (Fig. 11A, insets). The functional form of this dependence changes only slightly with the location of the recording site (Fig. 11B, dashed vs. solid line).

**Figure 11:**
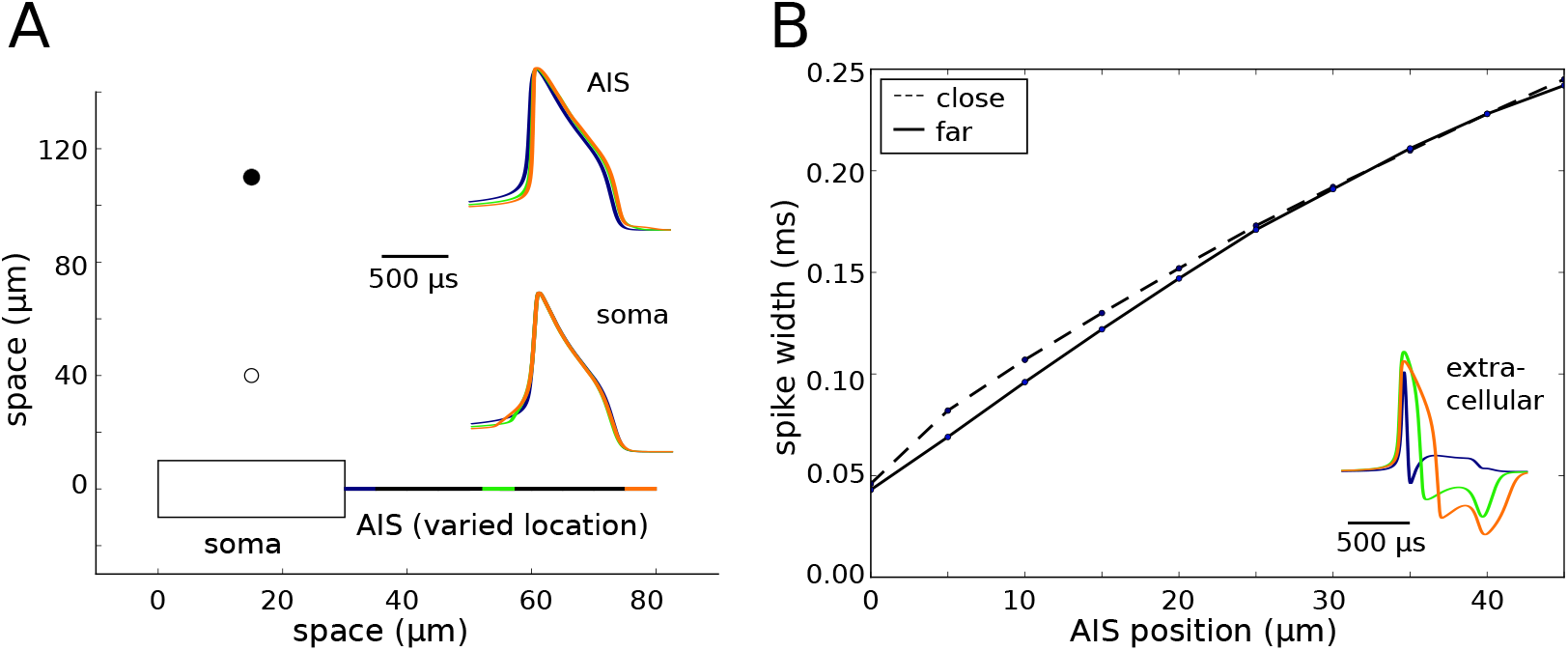
Width of the extracellular AP as a function of the soma–AIS separation. *A*: Schematic representation of the soma-axon model (bottom) and their relation to the recording points (dots above soma). The AIS position was systematically varied from 0 (directly attached to the soma) to 45 *μ*m. Insets: Waveforms of action potentials recorded intracellularly in the AIS (top inset) and the soma (bottom). The waveforms are normalized to the peak of somatic potential. *B*: Action potential width measured at half amplitude as a function of the AIS position for two different recording locations (close: 30 *μ*m from soma, far: 100 *μ*m from soma). Inset: Examples of extracellular AP waveshapes for 3 different locations of AIS (recorded 40 *μ*m above the soma).

## Discussion

Using detailed morphological models of reconstructed neurons and simplified soma-axon models we have shown that extracellular action potentials can be reconstructed from the current dipole formed by the soma and AIS at their initiation. We also show that the EAP shape depends on the position of the recording electrode with respect to the neuron promoting the extracellular contribution of different compartments of the neuron. In addition, while the width of the EAPs varies with the distance between the soma and the AIS, their amplitudes remain relatively constant.

The contribution of the AP to the extracellular field is shaped by the structure of the dendritic tree and the site of AP initiation. A large body of experimental data support the more distal initiation in the axon initial segment (Palmer and Stuart, 2006), but the impact of axonal initiation on the EAP had not been examined before. Using simplified models we showed that in the initial phase of the AP, the soma and AIS form a current dipole, whose contribution to the electric field decays inversely with the square of the distance from the dipole. At large distances (far-field approximation) the dipole contribution to the extracellular field does not depend on the separation between the AIS and the soma. In contrast, the width of the EAP increases with the soma/AIS separation. This soma-AIS dipole is different from the soma-dendrites dipole known from standard models (Gold et al., 2006). In fact, we showed that reducing the density of sodium channels in the AIS shifts AP initiation to the soma and as a consequence the extracellular potential is dominated by the soma-dendrites contribution.

Our results provide an important insight into the understanding of EAPs. It is known that the shape and the amplitude of the extracellular action potentials vary depending on the location of the recordings (Gold et al., 2006). Also, different types of neurons display extracellular action potentials of different width, such as excitatory cells, which tend to have broader extracellular action potentials when compared with interneurons (Barthó et al., 2004; McCormick et al., 1985), although there are exceptions (Vigneswaran et al., 2011). To separate action potentials of multiple neurons recorded extracellularly, it is common to use the waveform features of an extracellular action potential, such as the half-widths of the positive and negative peaks, the interval between them and the difference of their amplitudes (Lewicki, 1998; Einevoll et al., 2012). In addition, these and other waveform features sometimes allow the identification of neurons of different types (Peyrache et al., 2012; Dehghani et al., 2016). However, the significance of such features and their biophysical underpinnings are not completely understood. Numerical simulations of the extracellular field around reconstructed morphology of CA1 pyramidal neurons showed that the width of the extracellular action potential increases proportionally with the distance between the soma and the recording electrode (Gold et al., 2006). In addition, in this study the shape and amplitude of the extracellular potential was strongly affected by the channel densities in the dendrites and in the axon initial segment. In our work we show that the extracellular features of action potentials depend also on the exact location of their initiation site.

Finally, our results show that it should be possible, and of great interest, to follow experimentally the dynamic change of the AIS position by means of extracellular recordings. The length and distance of AIS from soma vary between neurons of same and different types (Fried et al., 2009; Kuba et al., 2006). Furthermore, the AIS is plastic and its length and position can change as a result of elevated activity which could occur due to plastic changes in a time scale of hours (Evans et al., 2015) to days (Grubb and Burrone, 2010; Evans et al., 2013; Muir and Kittler, 2014). This also happens as a consequence of a disease such as a stroke (Hinman et al., 2013; Schafer et al., 2009). Therefore, we expect that the shape of the EAP will vary according to the position of the AIS, such that long-term recordings from the same neuron could show gradual increase of the AP width. Since, the plasticity of AIS was never studied *in vivo* from intact neurons, this may open new methods of visualising such dynamic changes and investigating their functional role.

Our results are consistent with the large variability of EAP waveforms recorded in vivo (Fee et al., 1996; Harris et al., 2000). It is known that the waveshapes of the EAP depend on the position of the electrode, the morphology of the neuron and the densities of ion channels (Gold et al., 2007; Henze et al., 2000; Barthó et al., 2004; Pettersen and Einevoll, 2008). In particular, the presence of positive initial peak, as observed in our model, has been recognised in some studies (Palmer and Stuart, 2006). To further test our model experimentally, one could record the extracellular potential in vitro at multiple sites using multi-shank electrodes co-registered with the position of the soma and AIS. The AIS can be localised using fluorescent sodium channel markers (such as CoroNa) or immunostaining (for example, anykrin G is specific to AIS and nodes of Ranvier) (Zhou et al., 1998). This setup might allow for testing two new predictions of the model: 1) the presence of positive peak at the beginning of the EAP in the vicinity of soma-AIS region; 2) the width of the EAP as a function of the position of the AIS. In the latter case, we would need to visualise the change of AIS position dynamically probably over the course of many hours or days (Grubb et al., 2011). Such recordings are technically challenging, but are possible using present technology (Grubb and Burrone, 2010).

At the population level, the contribution of neurons to the local field potential (LFP) depends critically on the presence of voltage-dependent channels and neuronal morphology. For example, during the up state the LFP contains larger contributions from the active potassium and sodium currents than from synaptic currents (Reimann et al., 2013); similarly active conductances in the dendrites were shown to have major impact on the spectrum of the field potential (Ness et al., 2016). The structure of the dendritic tree has also been implicated in the generation of LFP signals (Lindén et al., 2010). Results in the present work suggest that the biophysics of the axon and the site of the action potential initiation may be additional factors determining the amplitude and the spectrum of the extracellular potential. The effects of the AIS position on LFP generated from a network of multi-compartmental model neurons is an interesting outlook of the present work.

## Acknowledgements

We thank Jennifer Goldman and Fabrizio Gabbiani for their comments on the manuscript. Research supported by the CNRS, the European Union (Human Brain Project H2020-720270), and the Agence Nationale de la Recherche (ANR-14-CE13-0003). MT was supported by Ecole des Neurosciences de Paris (ENP, http://www.paris-neuroscience.fr/) and DIM Cerveau et Pensée (http://dimcerveaupensee.fr/).

